# Body height in young adult men and risk of dementia later in adult life

**DOI:** 10.1101/748467

**Authors:** Terese Sara Høj Jørgensen, Gunhild Tidemann Christensen, Kaare Christensen, Thorkild IA Sørensen, Merete Osler

## Abstract

This study examined the relationship between body height and dementia and explored the impact of intelligence level, educational attainment, early life environment and familial factors. A total of 666,333 men, 70,608 brothers, and 7,388 twin brothers born 1939-1959 and examined at the conscript board were followed in Danish nationwide registers (1969-2016). Cox regression models were applied to analyze the association between body height and dementia. Within-brothers and within-twin pair analyses were conducted to explore the role of shared familial factors including partly shared genetics. In total, 10,599 men were diagnosed with dementia. The association between one z-score difference in body height and dementia (HR: 0.90, 95%CI: 0.89;0.90) was inverse and weakened slightly after adjustment for intelligence test scores and educational level. The within-brother analyses revealed a smaller estimate for dementia diagnosis the cohort analyses of brothers. The twin analysis showed similar, though less clear associations, which were not statistically significant.

## INTRODUCTION

Dementia poses substantial challenges for individuals and societies worldwide. ^1^ This has motivated studies of potential predictors and risk factors for dementia in recent years. ^1^ Development of dementia may be a result of both genetics and environmental exposures operating throughout the life course. ^2,3^ A newly published meta-analyses and systematic review showed that the risk of dementia may already be established early in life. ^4^ Body height has a strong genetic component and is at the same time influenced by environmental factors such as childhood diseases and nutrition. ^5,6^ Short height has been linked to development of dementia in a number of smaller studies (N=203-3,734) and to dementia as cause of death in a large study pooling 18 prospective cohorts (N=181,800). ^7–12^ Body growth may be related to dementia by being an indicator of brain and cognitive reserve and corresponding individual differences in the brain structure, which may imply differences in individual resilience toward development of dementia. ^4^ Another possible explanation of the height-dementia association is the correlation between body height and level of growth hormone that through hippocampal function and cognition has been linked to the risk of developing dementia. ^2,9,12^ Thus, rather than being a risk factors in itself, short body height is likely an indicator of harmful exposures early in life. ^7^ However, large scale high-quality longitudinal studies exploring the impact of early environmental factors and genetics to explain the link between body height and dementia are needed. ^2,4,7^ Previous studies question whether body height and dementia are solely linked by brain and cognitive reserve measured by intelligence and socioeconomic factors, or by shared underlying familial factors including genetics that may influence both body height and dementia. In this study, we examined whether body height in young adult men is associated with diagnosis of dementia while exploring whether intelligence test score, educational level, and underlying environmental and genetic factors shared by brothers explain the relationship.

## RESULTS

We followed 666,333 men for an average of 41.4 years from a baseline mean age of 22.1 years (due to delayed entry for some men) to a maximum age of 77 years. The mean age at measurement of height was 19.73 years (95%CI:19.73;19.74). Table 1 provides body height measures and information on dementia diagnoses in the three study populations of the total population of men. In total, 10,599 (1.6%) men were registered with dementia diagnosis during follow-up. The mean height was 176.8 cm (SD: 6.6). Table S2 shows that the mean height was greater among men with higher intelligence level, higher level of education, younger birth cohorts, and those living in the Capital region. Similarly, these groups also experiences fewer cases of dementia.

**Table 1.** Hazard ratios (HRs) and corresponding 95% confidence intervals (95%CI) of the association between taller body height at entry to adulthood and dementia diagnoses among the total population of men

| Descriptive statistics |  | HR (95% CI) for onset of dementia per One z-score higher |  |  |  |
| --- | --- | --- | --- | --- | --- |
| Mean height<br>(Standard deviation) | Dementia cases<br>Number (%) | Model 1<sup>*</sup> | Model 2a<sup>**</sup> | Model 2b<sup>***</sup> | Model 3<sup>****</sup> |
| 176.8<br>(6.6) | 10,599<br>(1.6) | 0.86<br>(0.85;0.87) | 0.88<br>(0.87;0.89) | 0.90<br>(0.89;0.90) | 0.90<br>(0.89;0.90) |

<sup>\*</sup>Model 1: Stratified by birthcohort and conscript board district. Age included as underlying scale of the model.
<sup>\*\*</sup>Model 2a: Model 1 + adjusted for educational level
<sup>\*\*\*</sup>Model 2b: Model 1 + adjusted for intelligence test scores
<sup>\*\*\*\*</sup>Model 3: Model 1 + adjusted for educational level and intelligence test scores

In Model 1, the estimated HR was 0.86 (95%CI:0.85-0.87), which attenuated slightly after the additional adjustments for educational level and intelligence test scores in Model 2a and 2b (Table 1). The fully adjusted estimates of the birth cohort-specific z-score of body height (Model 3) showed that for each one unit taller z-score of body height, the hazard ratio of dementia were 0.90 (95% CI:0.89;0.90). This was the same estimates as found for the model adjusted for intelligence test scores without educational level (Model 2b). Table S2 shows that the mean z-score is around 6.5 cm (range: 6.38-6.55) for all birth cohorts with small variations without any clear patterns of change over time. Figure 2 shows that the HR for the association between z-score of body height with dementia diagnosis is curve-linear; the HRs showed a slightly stronger increase with lower z-scores of body height below the reference of 0 than above the reference of 0.

Analyses of Model 3 with the risk period divided at age 60 years showed that the association between one z-score taller body height and dementia was smaller among men below (HR:0.87, 95%CI: 0.84;0.90) than above (HR:0.91, 95%CI: 0.90;0.92) 60 years (Table 2).

**Table 2.**
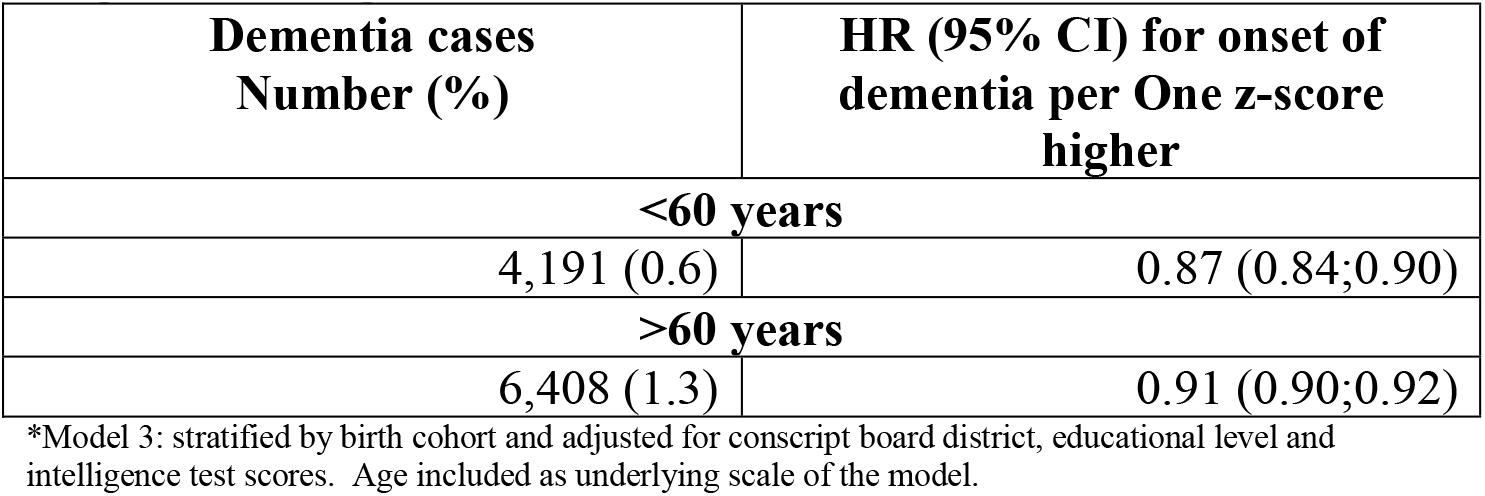
Fully adjusted* hazard ratios (HRs) of the association between taller body height at entry to adulthood and dementia diagnosis among all men

| **Dementia cases**<br>**Number (%)** | **HR (95% CI) for onset of**<br>**dementia per One z-score**<br>**higher** |
| --- | --- |
| **<60 years** |  |
| 4,191 (0.6) | 0.87 (0.84;0.90) |
| **>60 years** |  |
| 6,408 (1.3) | 0.91 (0.90;0.92) |

\*Model 3: stratified by birth cohort and adjusted for conscript board district, educational level and intelligence test scores. Age included as underlying scale of the model.

The Model 3 cohort analyses of brothers also showed that taller body height was associated with lower HR of dementia at the same level as in the full cohort analysis. The within-pair analyses showed lower point estimates, but with less precision as expressed by the wider confidence intervals (Table 3). For twins, the comparison of cohort and within-pair analyses revealed a pattern similar to that found for brothers, although the pattern was less clear and the estimates were statistically insignificant (Table 3).

**Table 3.** Fully adjusted* hazard ratios (HRs) and corresponding 95% confidence intervals (95%CI) of the association between taller body height at entry to adulthood and dementia diagnosis among brothers and twins

| **Descriptive statistics** |  | **HR (95% CI) for onset of dementia per one z-score higher** |  |
| --- | --- | --- | --- |
| **Mean height**<br>**(standard**<br>**deviation)** | **Dementia cases**<br>**Number (%)** | **Cohort analyses** | **Within-brother/twin analyses** |
| **Brothers** |  |  |  |
| 177.8 (6.6) | 597 (0.8) | 0.90 (0.82;0.98) | 0.78 (0.64;0.96) |
| **Twins** |  |  |  |
| 175.8 (6.7) | 107 (1.4) | 0.91 (0.73;1.16) | 0.86 (0.44;1.68) |

\*Model 3: Stratified by birth cohort and adjusted for conscript board district, educational level and intelligence test scores. Age included as underlying scale of the model.

## DISCUSSION

In this nationwide cohort of Danish men born from 1939 through 1959, we detected that taller body height at the entry to adulthood was associated with lower risk of dementia diagnosis later in life. The relationship attenuated, but persisted after adjustments for educational level and intelligence test scores. Within-brother analyses confirmed the findings of a relationship between taller body height and lower risk of dementia diagnosis, and even revealed a greater association than the cohort analyses. The findings from the within-twin analyses were in the same direction but with more uncertainty. These findings support a relationship between body height and dementia independent of the genetic relationship and of exposures to other shared early-life familial factors.

### Strengths and limitations

This study has a number of strengths. It is based on a very large unselected population followed from early adulthood until a maximum age of 77 years for dementia diagnoses. This study design is very important as previous findings may have underestimated the relationship due to selection of the healthiest part of the population, who are willing to participate, and have survived into old age. It is also a great advantage that we were able to investigate associations between body height and dementia adjusted for intelligence test scores and educational level. Intelligence together with educational level have been shown to be strong markers of cognitive reserve, which are hypothesized to be highly predictive of the age at which dementia is manifested and diagnosed. ^2^ We further included objective measures of body height rather than self-reported height with it’s implicit reporting errors. In the same line, this study included measures of body height from early adulthood, whereas previous studies have used measures from mid- and late-life. This could imply a bias if declining height since young adulthood ^13^ is more pronounced in socioeconomic dis-advantaged groups and those with lower later-life cognitive function. ^14^ Further, underdiagnoses of dementia has in some countries been shown to be greater in socioeconomically disadvantaged groups. ^15^ A final strength is that we were able to investigate the impact of familial factors, including partly shared genetics, by conducting within-brother and twin-pair analyses.

We acknowledge the limitations of the study. A potential concern is that even though dementia diagnoses in the Danish National Patient Register is likely to be correct, ^16,17^ under-diagnosis is possible. The registers and identifications to define dementia change over time (Supplementary Figure S1) and this difference in coverage could lead to differences in diagnostic validity and completeness at different time points during follow-up. However, we believe this is a minor issue as a previous study of the DCD data, which investigated the relationship between cognitive ability and dementia, only found small differences in risk estimates when using ICD8 and ICD10 codes, respectively, to identify dementia. ^18^ Another issue is that the recorded diagnosis of dementia has been shown to be less valid in younger populations ^16,17^ and our model tests indicated that the HR was slightly less pronounced at higher ages. We therefore conducted analyses where follow-up time was split at age 60 years. Both analyses of men below and above 60 years showed associations between taller body height and lower risk of dementia, but the association was more pronounced in the younger population. This may partly be explained by the less valid symptom profile in these patients and misdiagnoses of psychiatric or neurological conditions for dementia. ^17^ Furthermore, the role of familial factors cannot be fully explored. The analyses of brothers were incomplete by being restricted to mainly cover the birth cohorts 1953-59 and brothers with shared mothers, implying a mixture of twin brothers, full non-twin brothers and maternal half-brothers, but without paternal half-brothers. The specific contribution of genetic factors, independent of environmental factors, cannot be assessed by twin pair analyses without distinction between mono- and dizygotic twins. Finally, the generalizability of the findings to women is questionable. Previous findings on potential gender differences in the relationship between body height and dementia or poor cognition in old age are inconclusive, ^8,19^ but the incidence and prevalence of dementia are found to be lower in men than in women. ^20^

### Findings in light of previous studies

The findings of this current study provide substantial support to previous evidence of a link between body height and dementia. ^2,4,7–12^ All previous studies had accounted for educational level and other socioeconomic indicators, yet none of these studies had adjusted for intelligence level earlier in life. Intelligence level has been suggested to be a stronger marker of brain and cognitive reserve than educational level. ^2^ Intelligence level is furthermore correlated with body height ^21,22^ and by itself associated with dementia. ^18^.

In contrast to previous studies, we also investigated the impact of other potential early-life familial factors including genetics and socioeconomic resources in the family that may influence both body height and later risk of dementia. ^4^ Body height has been shown to have a strong genetic component with around 80% of the variation in populations being explained by genetic differences between individuals. ^5,23,24^ The genetic component of height has furthermore been found to be consistent across countries independent of living standards. ^5^

Interestingly, we found that the estimate for the relationship between body height and dementia was greater in the within-pair analyses than in the cohort analysis of brothers. Because the genetic variation in body height is smaller within brothers than between men in general, the results could be explained by genetic contribution to differences in body height between unrelated men that is not contributing to the differences in risk of dementia and thereby dilutes the association. Through this mechanism, the finding of a stronger association within brothers may, thus, be due to a less dilution of the effects of different harmful exposures early in life influencing both growth and risk of dementia. Thus, the findings that the link between body height and dementia is stronger when familial factors are adjusted for suggests that genetics has a minor role in the linkage between body height and dementia. However, it is important to acknowledge that the within-brother analyses were subject to greater uncertainty shown by wider confidence intervals. One previous Swedish study applied a twin design including 106 monozygotic twin pairs discordant for dementia, which indicated that the twin with the shortest height also more often was the one who developed dementia, supporting out interpretation, but the association provided insignificant estimates. ^10^ Two of the previous studies, which had adjusted for APOE genotype, being the strongest single genetic marker related to development of dementia, also supported that conclusion. In more detail, one of the studies detected an association between body height and dementia, ^8^ and the other study, examining length of extremities and dementia, found an association to arm length in both women and men, but only to knee height (i.e. distance from foot sole to the anterior surface of the thigh of the lower leg) in women and not in men. ^11^

In conclusion, taller body height at the entry to adulthood, supposed to be a marker of early-life environment, is associated with lower risk of dementia diagnosis later in life. The association persisted when adjusted for educational level and intelligence test scores, suggesting that height is not just acting as indicator of the cognitive reserve. Within-brother analyses confirm the findings of a relationship between taller height and dementia diagnosis, and suggest that the association may have common roots in early life environmental exposures that are independent of the family factors shared among brothers.

## METHODS

The study was based on the Danish Conscription Database (DCD), which, in brief, holds information registered at mandatory Danish conscription examinations in the years 1957 to 1984 for men born from 1939 through 1959. ^22^ All Danish men are requested by law to appear before the conscript board for a physical and mental examination between the ages of 18 and 26 years. We linked the data with the Danish Twin Register (DTR), the Danish Civil Registration System, the Danish National Patient Registry, the Danish Psychiatric Central Register, and the Danish National Prescription Register. ^25–29^ Figure 1 presents the selection of the three study populations; 1) men (N=666,333), 2) brothers (N=70,608), and 3) twins (N=7,388). Brothers and twins were identified by two different approaches. Brothers were identified from the population through maternal linkage in CPR and were mainly included from the birth-cohorts 1953-1959 where the linkage between mothers and offspring is complete in the register. Brothers were defined broadly by including all men with the same identified mother i.e. full-brothers, half-brothers, adopted brothers, and twins, triplets etc. Thus, the sample of brothers also include twins from these birth cohorts as they are registered with the same mother. However, for the twin population, twin pairs in all the birth cohorts 1939-1959 were identified by linking the DCD to the DTR. The project was evaluated and approved by the Danish Data Protection Agency: Jr nr 2014-41-2911. According to Danish law, ethical approval is not required for purely register-based studies.

**Figure 1.**
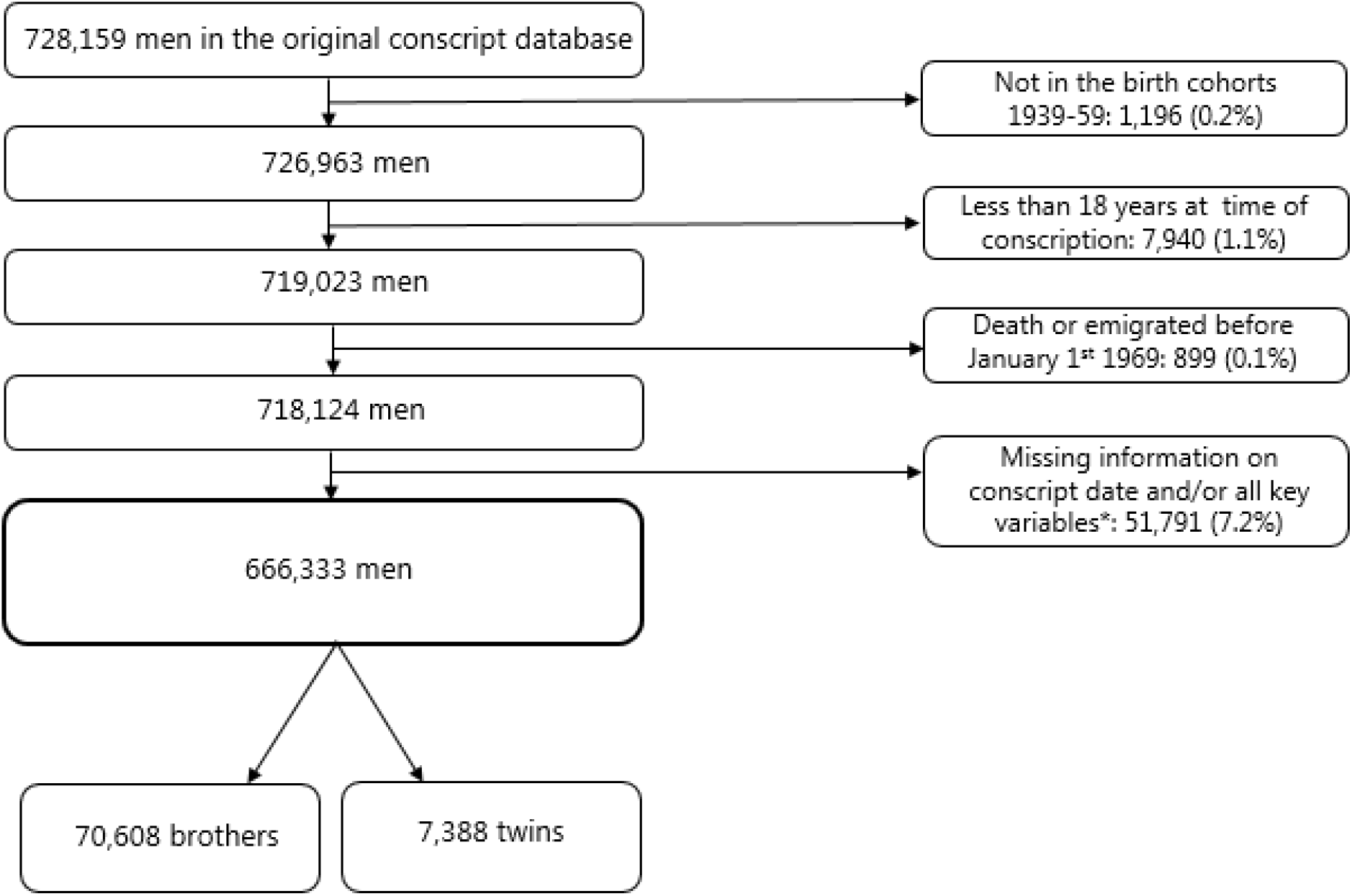
Selection of the study population

**Figure 2.**
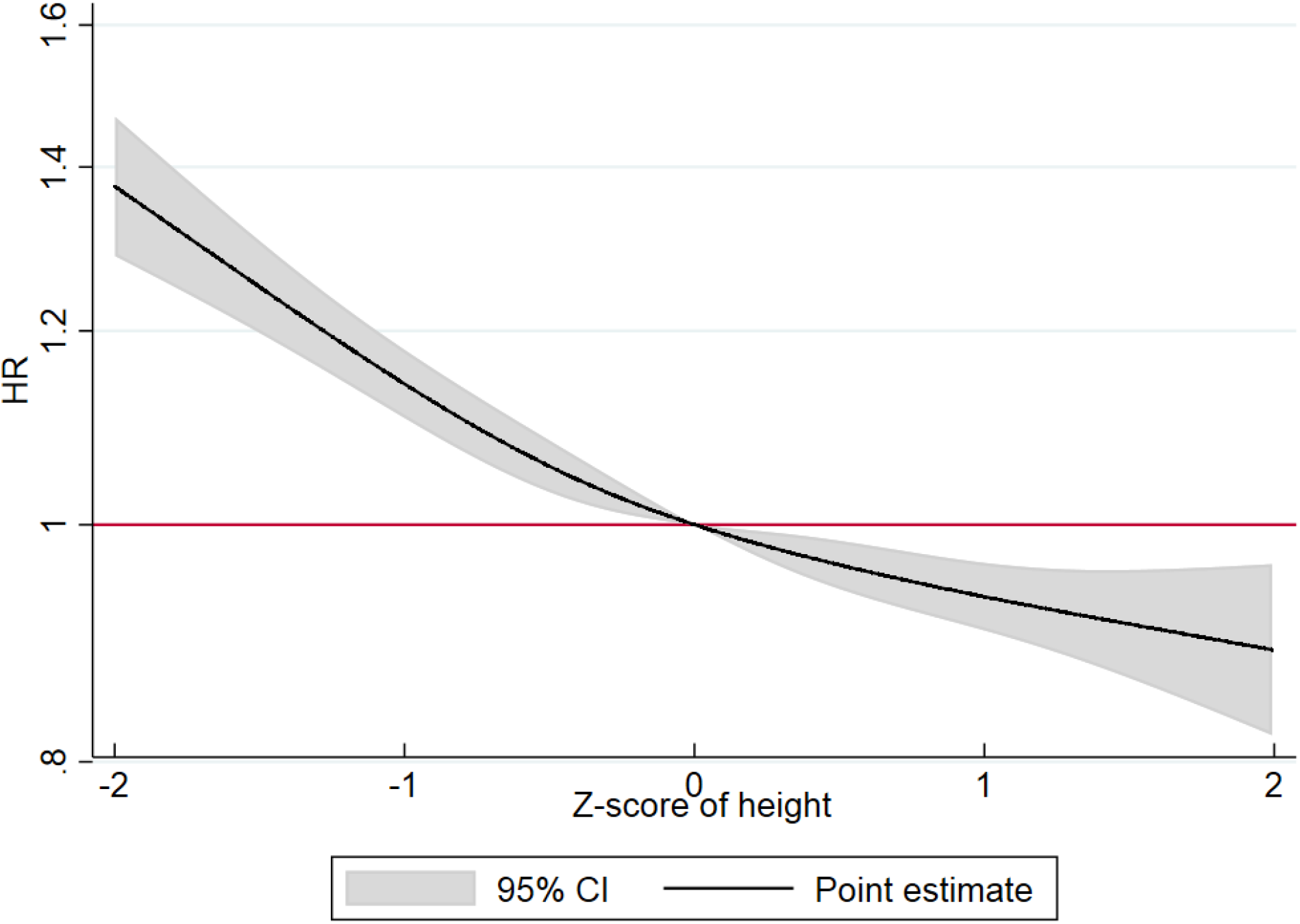
Hazard Ratios (HR) and corresponding 95% confidence intervals (95%CI) for the association between z-score of body height at then entry to adulthood as a cubic spline with 4 knots and dementia. A z-score of 0 is the reference value. Stratified by birth cohort and adjusted for conscript board district, educational level and intelligence test scores. Age included as underlying scale of the model. The analyses included the total population of men.

### Main variables

Body height at entry to adulthood was included as the main exposure in this study. Objective measures of body height without shoes at the time of conscript board examination was investigated as birth cohort specific z-scores (See supplementary Table S1). The z-scores were investigated as a continuous variable and as restricted cubic splines with four knots equally distributed over the range of the z-score variable: −1.63, −0.40, 0.36 and 1.65 with 0 as the reference.

Dementia diagnosis was measured by in- and out-patient contacts with international classification of disease (ICD) version 8 and ICD10 codes (ICD8: 290.00-290.99 and ICD10: F00.0-F03.9; G30.0-G30.9) in the Danish Psychiatric Central Register, since 1969, and in the Danish National Patient Registry since 1977. ^25,29^ Dementia diagnosis was furthermore identified by redeemed prescriptions of acetylcholinesterase inhibitor registered by Anatomic Therapeutical Chemical codes (N06D) in the Danish National Prescription Registry since 1995. ^28^ (See Figure S1 for a graphical illustration of the data collection time line for each of these registers). The dementia diagnoses in the Danish hospital registers have been shown to have a positive predictive value of 70% and 83% in two different randomly selected populations of 200 cases with dementia. The values vary between the two studies because of age differences in the study populations, the lowest positive predictive value was detected for the population with the youngest age group. ^16,17^

### Covariates

Intelligence test scores, educational level, age and district at conscript board examination, and birth cohorts were included as covariates in the analyses. Intelligence test scores were measured by the Børge Priens Prøve (BPP), which has been shown to be highly correlated with the full-scale Wechsler Adult Intelligence Scale score (R=0.82). ^30^ The score ranged from 0 to 78 and was categorized in deciles by 1) ≤21, 2) 22-27, 3) 28-31, 4) 32-35, 5) 36-38; 6) 39-42, 7) 43-45, 8) 46-49, 9) 50-53, 10) ≥54, and missing. Educational level at the time of examination was categorized by 1) short educational level: primary school, 2) medium educational level: vocational education and training as reference, 3) long educational level: secondary school, medium length education, high school, and academic educations, and 4) missing. Age was included as a continuous variable. Information about the eight conscript board examination districts was included as a categorical covariate with the following categories; 1) Copenhagen greater area incl. northern part of Zealand, 2) the remaining parts of Zealand and adjacent islands, 3) Funen and adjacent islands, 4) Mid and South parts of Jutland, 5) North and West Jutland, 6) Bornholm, 7) Southernmost part of Jutland, and 8) missing. The birth cohorts were included as a continuous variable ranging from 1939 through 1959.

### Statistical analyses

The association between body height at entry to adulthood and diagnosis of dementia was analyzed in three populations: 1) all men in the birth cohorts 1939-1959, 2) brothers mainly from the cohorts 1953-1959, and 3) twins in the birth cohorts 1939-1959.

We applied Cox proportional hazard regression models to estimate the associations between body height and subsequent dementia diagnosis (hazard ratios (HR) with 95% confidence intervals (CI)). For men with a conscript board examination date before January 1^st^ 1969, age at this date was used as baseline with a mean age of 22.1 years due to this delayed entry for some of the men. In contrast, men examined after January 1^st^ 1969 had baseline at the age of their conscript board examination. Men were included in the statistical analyses at baseline and followed until diagnosis of dementia, emigration, death, or end of follow-up (April 30^th^ 2016). Age was the underlying time scale of the model. Model assumptions of time-constant hazard ratios were tested by Schoenfeld residuals for continuous variables and log-minus-log curves for categorical variables. The model assumption was fulfilled for all variables. However, there was a tendency that the association between body height and dementia was stronger in the younger age groups, thus, a supplementary analyses with follow-up split at age 60 years were conducted to quantify this potential difference in the association between body height and dementia diagnosis before and after age 60 years.

To investigate the impact of adjustments for intelligence test scores and educational level as important explanatory factors, the analyses of the total population of men were conducted in four steps by Model 1 adjusted for conscript board examination district, Model 2a-b adjusted for, in addition to conscript board examination district, educational level and intelligence test scores, respectively, and finally, Model 3 adjusted for, in addition to conscript board examination district, both educational level and intelligence test scores. All the analyses were stratified by birth cohort to account for potential birth cohort effects and included a cluster term to account for the interdependence of observations between brothers. Finally, all models were by default adjusted for age as the underlying scale of the statistical models.

To explore the role of shared familial factors including environmental factors and partly shared genetics for the association between body height and dementia diagnosis, sibling populations (brothers and twins) were analyzed in a Model 3 standard cohort analyses and within-brother/twin pair analyses. In the cohort analyses each brother and twin was treated as an individual and a cluster term was included to account for the interdependence of observations. The within-pair analyses were stratified by a variable identifying the brother/twin pair cluster to make comparisons within groups of brothers and twin pairs. These analyses will provide estimates that, in addition to the included co-variates, are adjusted for shared family factors.

Statistical analyses were conducted in the statistical software packages Stata version 15.

## Acknowledgements

Authors thank D. Molbo, and E.L. Mortensen, who together with K. Christensen, M. Osler and T.I.A. Sørensen established the database. The work was supported by the Danish Medical Research Council [grant number 09-063599 and 09-069151], the Velux Foundation [grant number 31205], the Jascha Foundation, the Health Foundation (17-B-0033), Doctor Sofus Carl Emil Friis and Olga Doris Friis grant, and the Social Inequalities in Ageing (SIA) project, funded by NordForsk, project no. 74637.

## Supplementary

**Table S1.** Value of z-scores of body height in cm for each of the birth cohorts

|  | **Full study population** | **Brothers** | **Twins** |
| --- | --- | --- | --- |
| **Birth cohorts** | **Each z-score (cm)** |  |  |
| **1939** | 6.53 | 6.84 | 6.32 |
| **1940** | 6.45 | 6.35 | 6.62 |
| **1941** | 6.38 | 6.59 | 6.58 |
| **1942** | 6.42s | 6.31 | 6.36 |
| **1943** | 6.41 | 6.50 | 6.75 |
| **1944** | 6.45 | 6.61 | 6.43 |
| **1945** | 6.46 | 6.27 | 6.40 |
| **1946** | 6.47 | 6.47 | 6.35 |
| **1947** | 6.42 | 6.41 | 6.64 |
| **1948** | 6.48 | 6.49 | 6.67 |
| **1949** | 6.47 | 6.49 | 6.36 |
| **1950** | 6.44 | 6.39 | 6.60 |
| **1951** | 6.48 | 6.49 | 6.48 |
| **1952** | 6.50 | 6.53 | 6.40 |
| **1953** | 6.50 | 6.54 | 6.00 |
| **1954** | 6.55 | 6.64 | 6.98 |
| **1955** | 6.46 | 6.54 | 6.01 |
| **1956** | 6.52 | 6.55 | 6.85 |
| **1957** | 6.55 | 6.62 | 6.08 |
| **1958** | 6.50 | 6.52 | 6.88 |
| **1959** | 6.46 | 6.53 | 6.34 |

**Table S2.** Distribution of covariates for body height and dementia diagnoses among the total population of men

|  | **Body height**<br>**in cm** | **Dementia** |  |
| --- | --- | --- | --- |
|  | **Mean (SD)** | **No N, (%)** | **Yes N, (%)** |
| **Intelligence level (deciles)** |  |  |  |
| 1<sup>st</sup> | 174.0 (6.6) | 64,381 (97.3) | 1,783 (2.7) |
| 2<sup>nd</sup> | 175.1 (6.4) | 70,728 (98.0) | 1,476 (2.0) |
| 3<sup>rd</sup> | 175.6 (6.4) | 61,380 (98.2) | 1,101 (1.8) |
| 4<sup>th</sup> | 176.2 (6.4) | 70,687 (98.2) | 1,269 (1.8) |
| 5<sup>th</sup> | 176.6 (6.4) | 57,912 (98.6) | 841 (1.4) |
| 6<sup>th</sup> | 177.1 (6.4) | 80,365 (98.7) | 1,068 (1.3) |
| 7<sup>th</sup> | 177.7 (6.4) | 59,156 (98.6) | 831 (1.4) |
| 8<sup>th</sup> | 178.3 (6.4) | 70,967 (98.8) | 830 (1.2) |
| 9<sup>th</sup> | 178.7 (6.4) | 54,072 (98.8) | 648 (1.2) |
| 10<sup>th</sup> | 179.5 (6.3) | 60,682 (98.9) | 650 (1.1) |
| Missing | 177.0 (7.0) | 5,404 (98.2) | 102 (1.9) |
| **Educational level** |  |  |  |
| Short | 175.0 (6.5) | 165,994 (98.1) | 3,282 (1.9) |
| Medium | 175.9 (6.3) | 213,337 (98.2) | 3,952 (1.8) |
| Long | 178.7 (6.4) | 270,352 (98.8) | 3,247 (1.2) |
| Missing | 176.7 (6.9) | 6,051 (98.1) | 118 (1.9) |
| **Birth cohorts** |  |  |  |
| 1939-1944 | 175.2 (6.4) | 171,911 (97.0) | 5,338 (3.0) |
| 1945-1949 | 176.3 (6.5) | 151,927 (98.3) | 2,580 (1.7) |
| 1949-1954 | 177.4 (6.5) | 191,285 (99.0) | 1,903 (1.0) |
| 1955-1959 | 178.7 (6.5) | 140,611 (99.5) | 778 (0.6) |
| **Conscript board district** |  |  |  |
| Copenhagen greater area incl. northern part of Zealand | 177.8 (6.7) | 202,091 (98.1) | 3,829 (1.9) |
| The remaining parts of Zealand and adjacent islands | 176.0 (6.6) | 94,369 (98.4) | 1,554 (1.6) |
| Funen and adjacent islands | 176.3 (6.5) | 54,728 (98.2) | 1,032 (1.8) |
| Southern and mid-eastern parts of Jutland | 176.9 (6.5) | 117,831 (98.8) | 1,472 (1.2) |
| North and West Jutland | 176.4 (6.4) | 115,775 (98.8) | 1,452 (1.2) |
| Bornholm | 175.3 (6.5) | 6,993 (98.4) | 114 (1.6) |
| The reunited southern part of Jutland | 176.4 (6.5) | 62,123 (98.2) | 1,112 (1.8) |
| Missing | 175.6 (6.2) | 1,824 (98.2) | 34 (1.8) |

**Figure S1.**
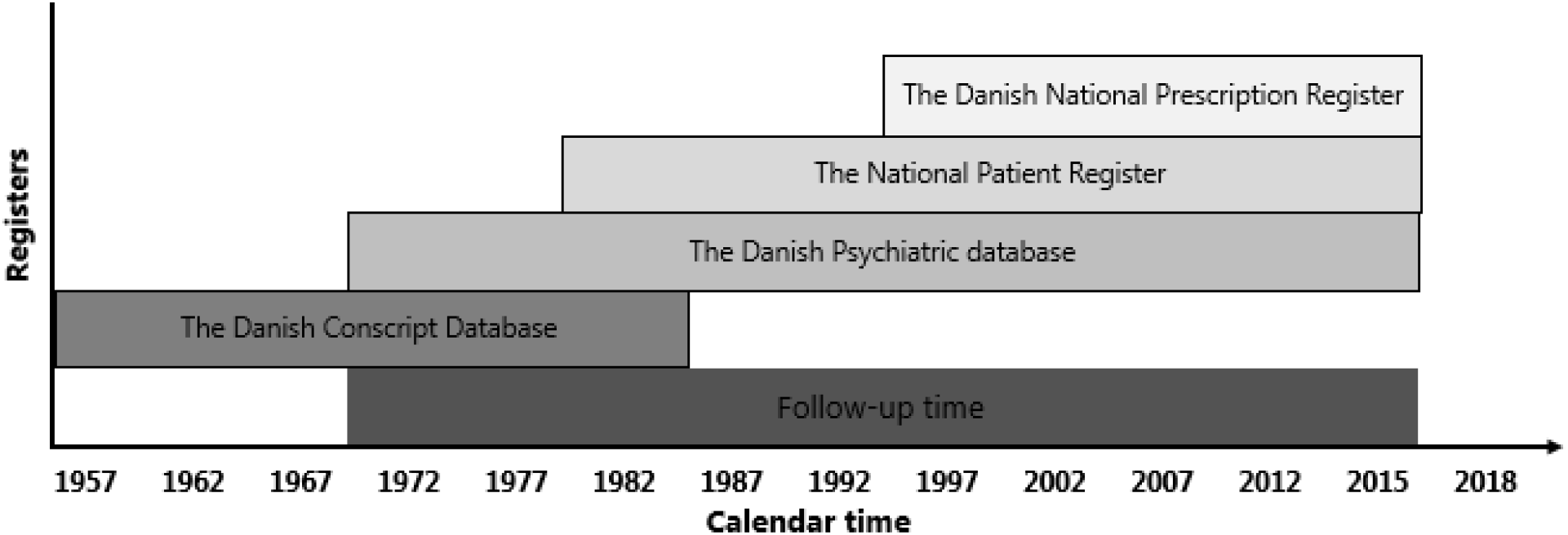
Time line of data collection from each of the registers and the total follow-up time

## REFERENCES

1. Livingston G, Sommerlad A, Orgeta V, et al. Dementia prevention, intervention, and care. The Lancet 2017;390(10113): 2673–734.

2. Borenstein AR, Copenhaver CI, Mortimer JA. Early-life risk factors for Alzheimer disease. Alzheimer Dis Assoc Disord 2006;20(1): 63–72. doi: 10.1097/01.wad.0000201854.62116.d7.

3. Bird TD. Genetic Factors in Alzheimer’s Disease. New England Journal of Medicine 2005;352(9): 862–4.

4. Wang XJ, Xu W, Li JQ, Cao XP, Tan L, Yu JT. Early-Life Risk Factors for Dementia and Cognitive Impairment in Later Life: A Systematic Review and Meta-Analysis. J Alzheimers Dis 2019;67(1): 221–9. doi: 10.3233/JAD-180856.

5. Jelenkovic A, Hur YM, Sund R, et al. Genetic and environmental influences on adult human height across birth cohorts from 1886 to 1994. Elife 2016; 5.(doi): 10.7554/eLife.20320.

6. Jelenkovic A, Sund R, Hur YM, et al. Genetic and environmental influences on height from infancy to early adulthood: An individual-based pooled analysis of 45 twin cohorts. Sci Rep 2016;6:28496.(doi): 10.1038/srep28496.

7. Russ TC, Kivimaki M, Starr JM, Stamatakis E, Batty GD. Height in relation to dementia death: individual participant meta-analysis of 18 UK prospective cohort studies. Br J Psychiatry 2014;205(5): 348–54. doi: 10.1192/bjp.bp.113.142984.

8. Petot GJ, Vega U, Traore F, et al. Height and Alzheimer’s disease: findings from a case-control study. J Alzheimers Dis 2007;11(3): 337–41.

9. Abbott RD, White LR, Ross GW, et al. Height as a marker of childhood development and late-life cognitive function: the Honolulu-Asia Aging Study. Pediatrics 1998;102(3 Pt 1): 602–9. doi: 10.1542/peds.102.3.602.

10. Gatz M, Mortimer JA, Fratiglioni L, et al. Potentially modifiable risk factors for dementia in identical twins. Alzheimers Dement 2006;2(2): 110–7. doi: 10.1016/j.jalz.2006.01.002.

11. Huang TL, Carlson MC, Fitzpatrick AL, Kuller LH, Fried LP, Zandi PP. Knee height and arm span: a reflection of early life environment and risk of dementia. Neurology 2008;70(19 Pt 2): 1818–26. doi: 10.212/01.wnl.0000311444.20490.98.

12. Beeri MS, Davidson M, Silverman JM, Noy S, Schmeidler J, Goldbourt U. Relationship between body height and dementia. Am J Geriatr Psychiatry 2005;13(2): 116–23. doi: 10.1176/appi.ajgp.13.2.116.

13. Cline MG, Meredith KE, Boyer JT, Burrows B. Decline of height with age in adults in a general population sample: estimating maximum height and distinguishing birth cohort effects from actual loss of stature with aging. Hum Biol 1989;61(3): 415–25.

14. Huang W, Lei X, Ridder G, Strauss J, Zhao Y. Health, Height, Height Shrinkage, and SES at Older Ages: Evidence from China. American economic journal Applied economics 2013;5(2): 86–121.

15. Lang L, Clifford A, Wei L, et al. Prevalence and determinants of undetected dementia in the community: a systematic literature review and a meta-analysis. BMJ Open 2017;7(2): e011146–e.

16. Phung TK, Andersen BB, Hogh P, Kessing LV, Mortensen PB, Waldemar G. Validity of dementia diagnoses in the Danish hospital registers. Dement Geriatr Cogn Disord 2007;24(3): 220–8. doi: 10.1159/000107084. Epub 2007 Aug 10.

17. Salem LC, Andersen BB, Nielsen TR, et al. Overdiagnosis of dementia in young patients - a nationwide register-based study. Dement Geriatr Cogn Disord 2012;34(5-6): 292–9. doi: 10.1159/000345485. Epub 2012 Nov 30.

18. Osler M, Christensen GT, Garde E, Mortensen EL, Christensen K. Cognitive ability in young adulthood and risk of dementia in a cohort of Danish men, brothers, and twins. Alzheimers Dement 2017;13(12): 1355–63. doi: 10.016/j.jalz.2017.04.003. Epub May 19.

19. West RK, Ravona-Springer R, Heymann A, et al. Shorter adult height is associated with poorer cognitive performance in elderly men with type II diabetes. J Alzheimers Dis 2015;44(3): 927–35. doi: 10.3233/JAD-142049.

20. Matthews FE, Stephan BC, Robinson L, et al. A two decade dementia incidence comparison from the Cognitive Function and Ageing Studies I and II. Nat Commun 2016;7:11398. (doi): 10.1038/ncomms11398.

21. Beauchamp JP, Cesarini D, Johannesson M, Lindqvist E, Apicella C. On the sources of the height-intelligence correlation: new insights from a bivariate ACE model with assortative mating. Behavior genetics 2011;41(2): 242–52.

22. Christensen GT, Molbo D, Angquist LH, et al. Cohort Profile: The Danish Conscription Database(DCD): A cohort of 728,160 men born from 1939 through 1959. 2014;(1464-3685 (Electronic)).

23. Lango Allen H, Estrada K, Lettre G, et al. Hundreds of variants clustered in genomic loci and biological pathways affect human height. Nature 2010;467: 832.

24. McEvoy BP, Visscher PM. Genetics of human height. Econ Hum Biol 2009;7(3): 294–306. doi: 10.1016/j.ehb.2009.09.005. Epub Sep 17.

25. Mors O, Perto GP, Mortensen PB. The Danish Psychiatric Central Research Register. Scand J Public Health 2011;39(7 Suppl): 54–7. doi: 10.1177/1403494810395825.

26. Skytthe A, Kyvik KO, Holm NV, Christensen K. The Danish Twin Registry. Scand J Public Health 2011;39(7 Suppl): 75–8. doi: 10.1177/1403494810387966.

27. Pedersen CB. The Danish Civil Registration System. Scand J Public Health 2011;39(7 Suppl): 22–5. doi: 10.1177/1403494810387965.

28. Kildemoes HW, Sorensen HT, Hallas J. The Danish National Prescription Registry. Scand J Public Health 2011;39(7 Suppl): 38–41. doi: 10.1177/1403494810394717.

29. Lynge E, Sandegaard JL, Rebolj M. The Danish National Patient Register. Scand J Public Health 2011;39(7 Suppl): 30–3. doi: 10.1177/1403494811401482.

30. Mortensen E, M. Reinisch J, Teasdale T. Intelligence as Measured by the WAIS and a Military Draft Board Group Test; 1989.

